# Root System Architecture Reorganization Under Decreasing Soil Phosphorus Lowers Root System Conductance of *Zea mays*

**DOI:** 10.1101/2024.05.31.596894

**Authors:** Felix Maximilian Bauer, Dirk Norbert Baker, Mona Giraud, Juan Carlos Baca Cabrera, Jan Vanderborght, Guillaume Lobet, Andrea Schnepf

**Author notes:** **Corresponding author:** Felix Maximilian Bauer.

## Abstract

The global supply of phosphorus is decreasing. At the same time, climate change reduces the water availability in most regions of the world. Insights on how decreasing phosphorus availability influences plant architecture is crucial to understand its influence on plant functional properties, such as the root system’s water uptake capacity. In this study we investigated the structural and functional responses of *Zea mays* to varying phosphorus fertilization levels focusing especially on the root system’s conductance. A rhizotron experiment with soils ranging from severe phosphorus deficiency to sufficiency was conducted. We measured architectural parameters of the whole plant and combined them with root hydraulic properties to simulate time-dependent root system conductance of growing plants under different phosphorus levels. We observed changes of the root system architecture, characterized by decreasing crown root elongation and reduced axial root radii with declining phosphorus availability. Modeling revealed that only plants with optimal phosphorus availability sustained a high root system conductance, while all other phosphorus levels led to a significantly lower root system conductance, both under light and severe phosphorus deficiency. We postulate that phosphorus deficiency initially enhances root system function for drought mitigation but eventually reduce biomass and impairs root development and water uptake in prolonged or severe cases of drought. Our results also highlight the fact that root system organization, rather than its total size, is critical to estimate important root functions.

## Introduction

The exploitation of finite natural resources poses new challenges to agriculture. The supply of phosphorus (P), a vital nutrient derived from finite resources, will decrease (Marschner, 2011). P scarcity has been identified as a likely key factor limiting future food availability, emphasizing the urgency of addressing this issue (Cordell et al., 2009). The predicted time for “peak phosphorus”, i.e. the time at which global P production reaches its maximum due to the depletion of reserves and declines again immediately afterwards, is estimated around the early to mid-21st century (Reijnders, 2014). Additionally, excessive use of P fertilizer significantly impacts the environment by contributing to eutrophication, which is harming open water bodies and leads to aquatic plant and algae growth, impairing water quality for other organisms and limiting water use for drinking, recreation, and industry (Randall, 2003). Especially in lakes, rivers, estuaries, and coastal oceans over-enrichment with P is a widespread problem (Carpenter et al., 1998). Most of the P stored in water bodies originates from agricultural and urban activities. P fertilizers dissolve quickly, releasing P faster than plants can absorb it, thus these fertilizers are highly prone to being lost by erosion. P that is not used by plants or object to runoff losses is immobilised in the soil and subsequently not available for plants anymore. For these reasons, a reduction of P fertilization is required.

Concurrently to the impending shortage of P, climate change is anticipated to lead to a scarcity of water across various regions around the globe (Gosling and Arnell, 2013). In view of this future water shortage it is therefore crucial to gain an advanced understanding of how the reduced availability of P affects the plant’s architecture, specifically functional alterations, related to the plant’s capacity for water uptake through its root system (Fry et al., 2018).

*Zea mays* is one of the most important crops world wide and crucial for human nutrition (Ranum et al., 2014). Maize is sensitive to P deficiency and in at least 30% of the cultivated land area with *Zea* cultivars, P is limiting (Heuer et al., 2017). Especially for *Zea mays* it is known that canopy development is inhibited by P deficiency, leading to yield decline. The plant’s architecture changes under P limitation. P deficiency is often associated with reduced growth and rigid appearance of shoots. Limited P availability also induces changes in root architecture. Studies report different morphological changes, such as the inhibitions of primary root growth, shallower axial root angle, or various changes in lateral root growth, such as the reduction of lateral root growth in the field, but also an increase in lateral branching in plants with few axial roots (zero order roots (Borch et al., 1999; Zhu and Lynch, 2004; Marschner, 2011), often resulting in a higher root to shoot biomass ratio (Lynch et al., 2005). Furthermore, an increase in crown root number has been reported to be beneficial under P deficiency (Sun et al., 2018). A reduced root radii was described as a response of *Zea mays* to reduced P availability in soil (Sheng et al., 2012; Zhang et al., 2012). However, no direct functional relationship between soil P availability and this responses has yet been established. Additionally, under field conditions, most plant responses are measured in rather coarse metrics and do not provide direct response functions (Lopez et al., 2023). Although a variety of different plant responses were reported, it remains uncertain which parameters (non-aggregated, directly measurable attributes, such as type depended root length and number) have an direct impact on aggregated structural and functional root system traits, such as total root system volume or plants water uptake capacity.

Main drivers for the plant’s water uptake capacity are the root system architecture and anatomy (Steudle, 2001). Root plasticity refers to the ability of plant roots to alter their growth depending on environmental conditions. Root architecture depends on the root system’s plasticity in the soil. Root anatomy relates to the internal structure of the root. Together, they govern the root hydraulic properties. The root hydraulic properties are essential to transport water from the soil to the roots and the to the aboveground organs. Radial hydraulic conductivity (*k_r_*) is a measure of a root’s ability to take up water from the soil into its vascular system. In contrast, axial hydraulic conductance (*K_x_*) relates to the efficiency of water transport along the length of the root’s main axis (Steudle, 2001). Changes in these properties at the cellular and organ levels can impact the root’s overall hydraulic function that might alter the root system conductance (*K_rs_*). A change in *K_rs_* affects the plant’s ability to uptake water (Meunier et al., 2020). Generally, the variability in *K_rs_* can be very high. The variation in root anatomy is especially strong among *Zea* lines (Rishmawi et al., 2023). A lot of different environmental influences, such as drought or salinity, lower K_rs_ (Aroca et al., 2011). Also the root system age has an important influences on *K_rs_* (Baca Cabrera et al., 2024). P deficiency was also suggested as an influencing factor for lowering *K_rs_* in different species (Shanguan et al., 2005; Mu et al., 2006; Li et al., 2009). The relationship between P availability and key architectural root system parameters that drive changes in *K_rs_* is not well understood. Moreover, *Zea mays* was rarely the object of studies investigating the influences of P deficiency on *K_rs_*. However, with experimental setups, it is especially challenging to quantify solely the effects of P limitation on whole crop and canopy development and its consequences on relevant physiological processes, such as water uptake related functions. Especially for *k_r_*, experimental measurements require complex setups, such as root pressure probe (Frensch and Steudle, 1989), measuring water flow of pruned roots within a pressure chamber (Zwieniecki et al., 2002), or use the high pressure flow meter device on whole root systems for root system conductance, as proposed by (Tyree et al., 1994). The necessity of measuring *k_r_* at several locations, in case of its variation along the root axis or for various root types, makes its experimental evaluation more challenging. Inverse modeling is a newer, additional method to obtain *k_r_* and *K_x_* values (Couvreur et al., 2018).

Functional-structural plant models (FSPMs) are a suitable tool to help investigate and interpret the reaction of the plant to a changing environment, such as the absence of a crucial nutrient. They can bridge the gap between the sub-organ and whole plant level, and thus simulate mechanistically emerging plant phenotypes caused by the interaction of processes at smaller scales, such as the effect of radial and axial water fluxes through root segments on the whole plant water uptake. Indeed, FSPMs are computational frameworks that simulate plant growth by integrating physiological functions with (3D-)structural representations of plant organs. In the context of P, FSPMs have already been used to test hypotheses regarding changing root architecture of *Zea mays*, such as a bigger inter-lateral distance (Postma et al., 2014) and a higher amount of seminal roots regarding its advantages for P uptake (Perkins and Lynch, 2021).

Although it has already been shown that root architecture and shoot size adaptation are affected by soil P deficiency, transferring these findings directly to the sub-organ level is very complex without more detailed experimental investigation of how architecture changes at a high spatio- temporal resolution. Previous studies had however a coarse temporal or spatial resolution or focused on specific organs (Sun et al., 2018). Moreover, it is also suggested that potential reactions to P deficiency can already occur in very early growth stages (Brunel-Muguet et al., 2014), whereas experimental studies focused on older plants (Pereira et al., 2020). To the best of our knowledge there are currently no studies investigating the whole plant’s architectural response of *Zea mays* to different levels of phosphorus availability, including the influence of this response and its consequences for *K_rs_*.

This work aims at understanding which root and shoot architectural parameter are responding to four decreasing P levels from sufficient to severe deficient and how this will affect the plants root system capacity for water uptake. Therefore, this study has two main objectives:

1. Identify experimentally which structural parameters of maize organs show the strongest responses to soil phosphorus availability.
2. Parameterize and use *Zea mays* FSPMs from experimental data of to analyze how root system conductance in maize adapts to the different P availability levels.

## Results

### Plant structural responses to soil P level

The influences of P deficiency on the architecture of young root systems appears complex. Although we observed a reorganisation in a lot of different architectural root system traits, the clearest significant trends in root trait responses to P deficiency can be seen in the radius of axial roots and the elongation rate of crown roots (Figure 1A&B). The radii of axial roots significantly increased with the amount of P fertilized. Only for the initial leaf elongation rate we found a significant architectural response of the shoot to P availability. The initial elongation rate was significantly higher for the highest P level (P3) compared to the two lowest P levels (P0-P1) (Figure 1C). Consequently, the maximal leaf area showed an increasing trend as well. Although stem length and diameter also increased slightly with higher P supply, the differences between the P levels were not significant. The destructively measured root to shoot biomass ratios show a decreasing trend with increasing P availability (Figure 1D). We performed a principal component analysis (PCA), including the significant response parameters to P deficiency, which clearly demarcates clusters for each phosphorus treatment level, with minimal overlap between the confidence ellipses. This suggests a strong grouping effect in our data, reflective of the distinct phosphorus treatments applied (Figure 2). The PCA further revealed that axial root radii are closely associated with soil P content, while crown root elongation showed a notable correlation with the P to dry matter ratio (*PB*, *mg P g biomass*^−1^). A similar positive correlation was observed between soil P and leaf elongation rate. Therefore, we considered crown root elongation rate and axial root radii as plastic response parameters for root system changes. Additionally, the leaf elongation rate was considered as shoot response to P availability.

**Figure 1:**
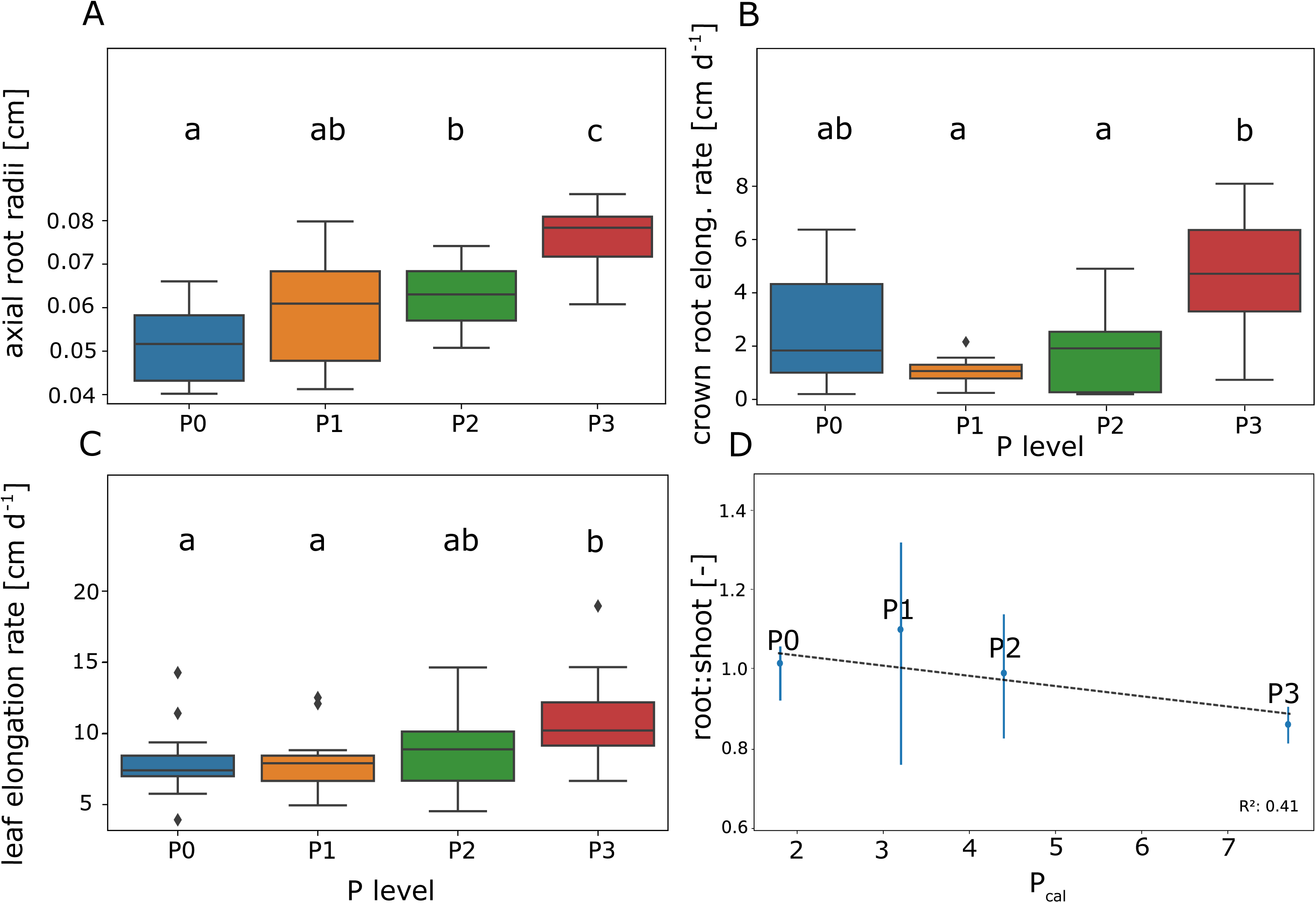
A: Axial root radii, B: Initial crown root elongation rate, C: Initial elongation rate of the leaves. D: Root:Shoot ratio for different soil P availability levels. *p<0.05*

**Figure 2:**
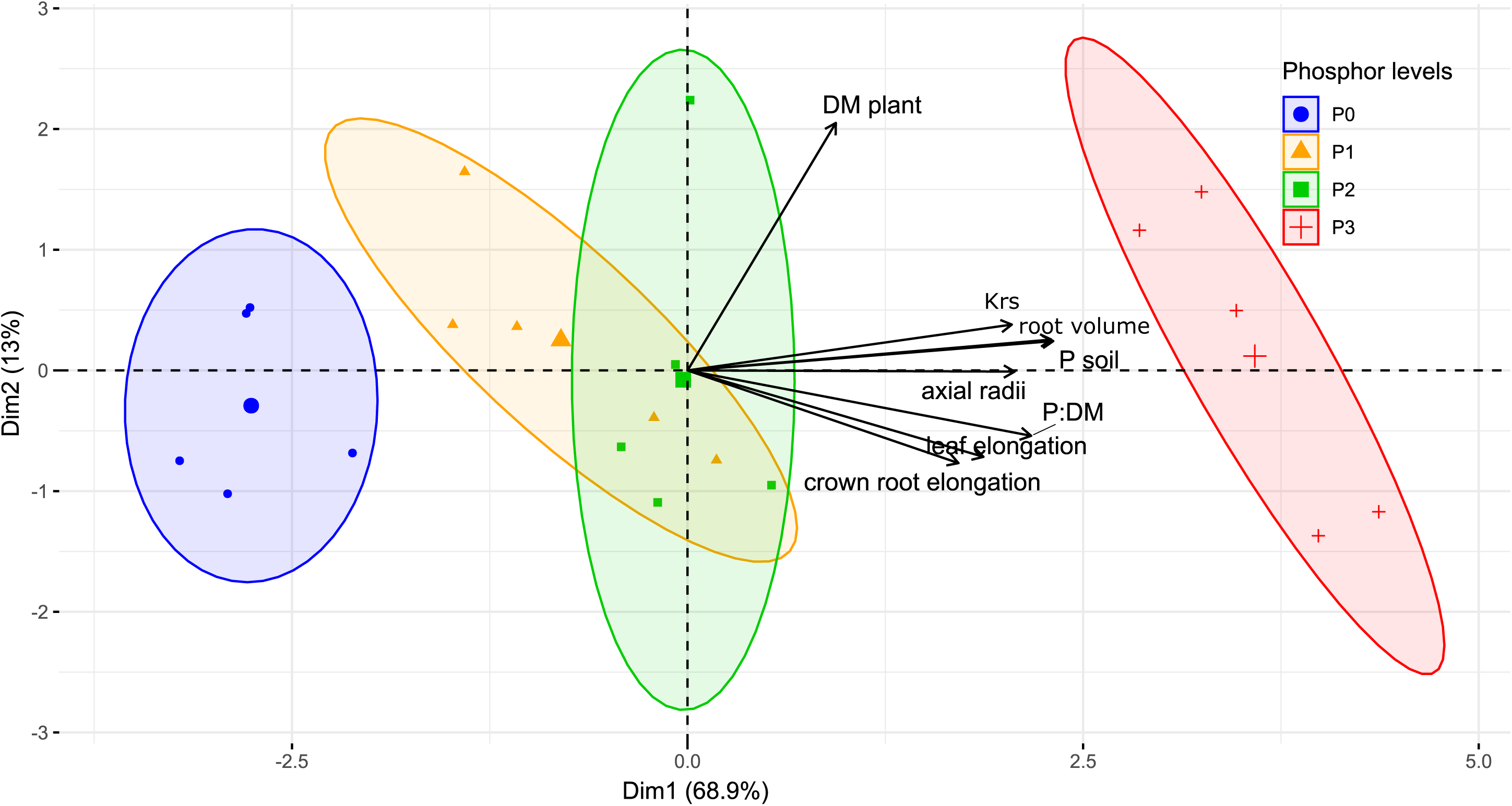
Principal component analysis (PCA) to identify the contribution of the plant parameters to the response of *Zea mays* to P deficiency. The big symbols corresponds to the centroids for the different P treatments.

For the radii of axial roots (*a*_*ax*_) we found a direct linear relationship to P available in soil (Figure 3A) within the measured upper boundary ( *P*_*max*_) and lower boundary *P*_*min*_(*mg hg soil* (100 *g*^−1^) of soil P, described as eq. 1.

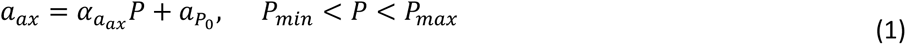

where parameter *α*_*aax*_ defines the increase in radii per unit of P in soil and *a*_*P*0_ the intercept of the response that represents the radii at the theoretical situation of no available P in soil. We found that the crown root elongation rate *a*_*rc*_ (*cm d*^−1^) is a response to the ratio of *P*_*soil*_ to dry matter (DM, *g*) of the plant (eq. 2).

**Figure 3:**
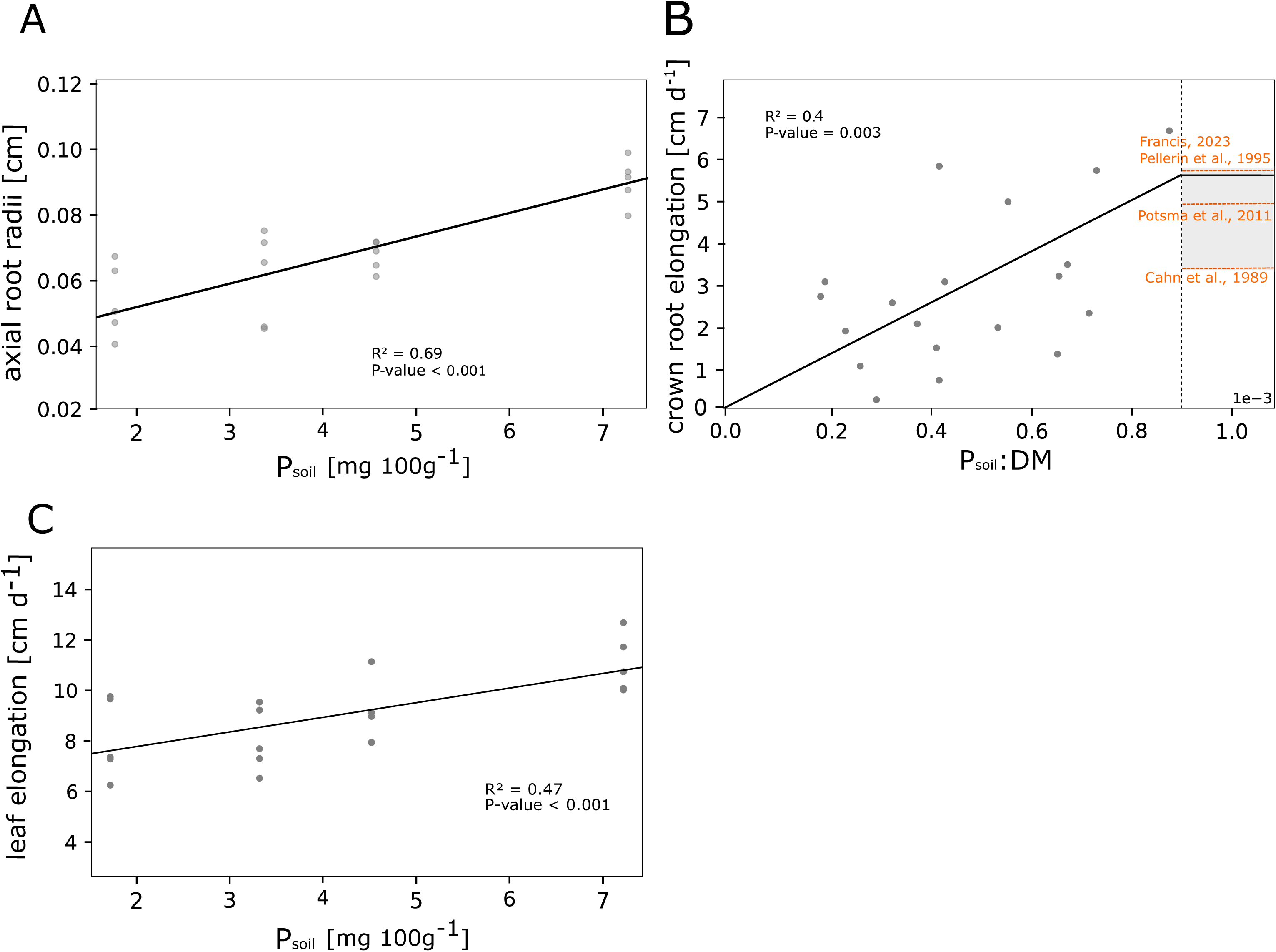
Response of axial roots radii (A), crown root elongation (B) and leaf elongation rate (C) to different soil P availability levels.

**Figure 4:**
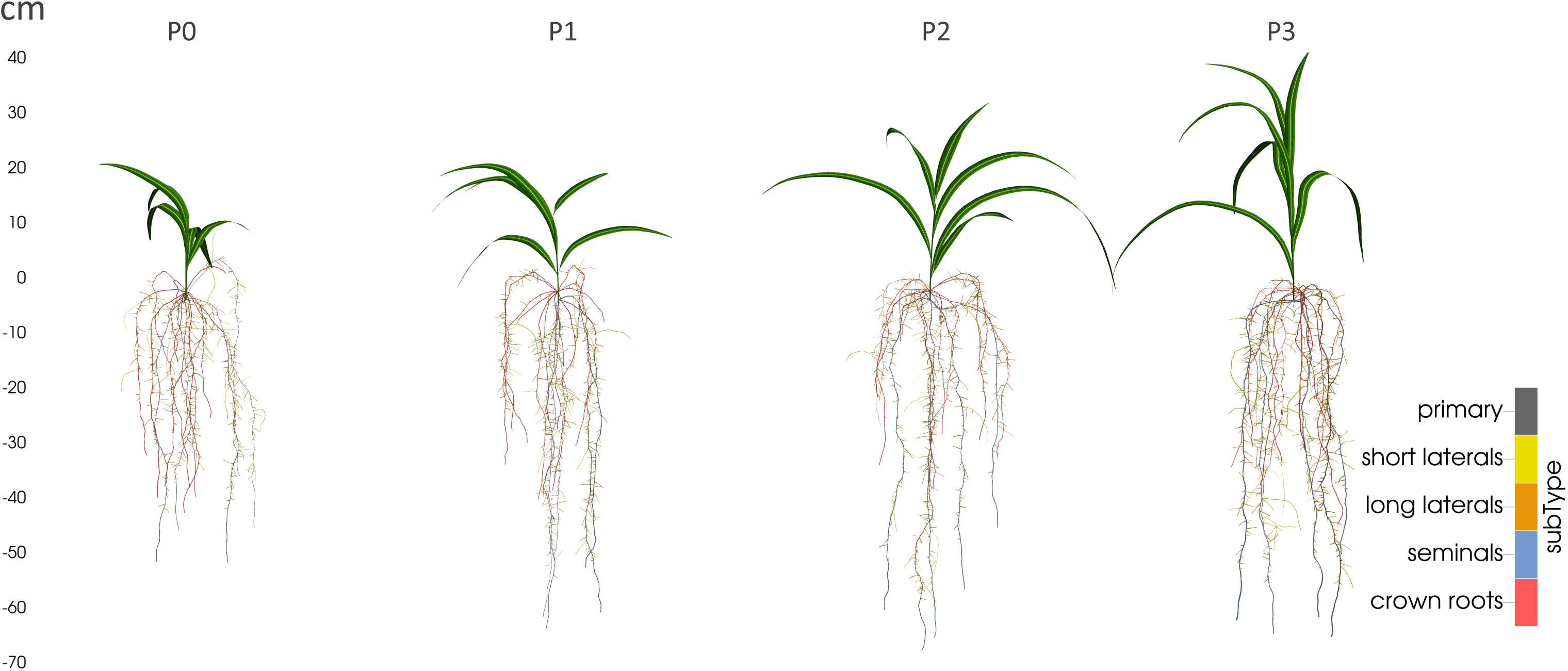
Simulated plant structure with CPlantBox for all P levels. The given subtypes correspond to the denomination of the root types.

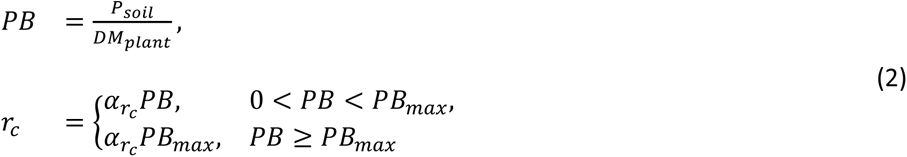

where *α*_*rc*_ is the increase in elongation per unit *PB*. *PB*_*max*_ describes the maximal *PB* we measured, which however aligns with several maximal crown root elongation rates, measured by other studies (Figure 3B).

The initial leaf elongation rate (*r*_*l*_, *cm d*^−1^) is a linear function of the P available in soil and described by eq. 3 (Figure 3C).

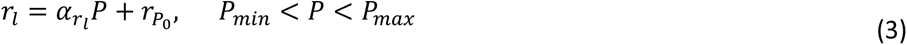

where *α*_*rl*_ (*cm d*^−1^) is the increase in elongation per unit P, while *r*_*P_0_*_ (*cm mg P d*^−1^ *hg soil*) is the intercept at the theoretical situation of no soil P. *P*_*min*_ and *P*_*max*_ (*mg hg soil*) describe the lower and upper boundaries of *P* for the *r*_*l*_ variations. Our observations revealed that the leaf area was maintained for plants with higher P supply and sharply decreased at the two lowest P levels (Table 3). The root volume increased linearly with the amount of available soil P (Supplementary Figure 1).

For every P level, a complete CPlantBox parameter-set was created. A full list of the parameters, including root system initializing parameters, as well as root and shoot specific parameters can be found in Tables 1, 3 and 3, respectively. We moreover created a FSPM, which is simulating dynamic growth of *Zea mays* cv. B73 under different soil P levels and modified only the identified key parameters (see section “P levels strongly influence axial root radius and crown root elongation”) according to the measured P levels. We compared the time dependent simulated total root system volume and found no relevant absolute differences between the same treatments (Supplementary Figure 1).

**Table 1:**
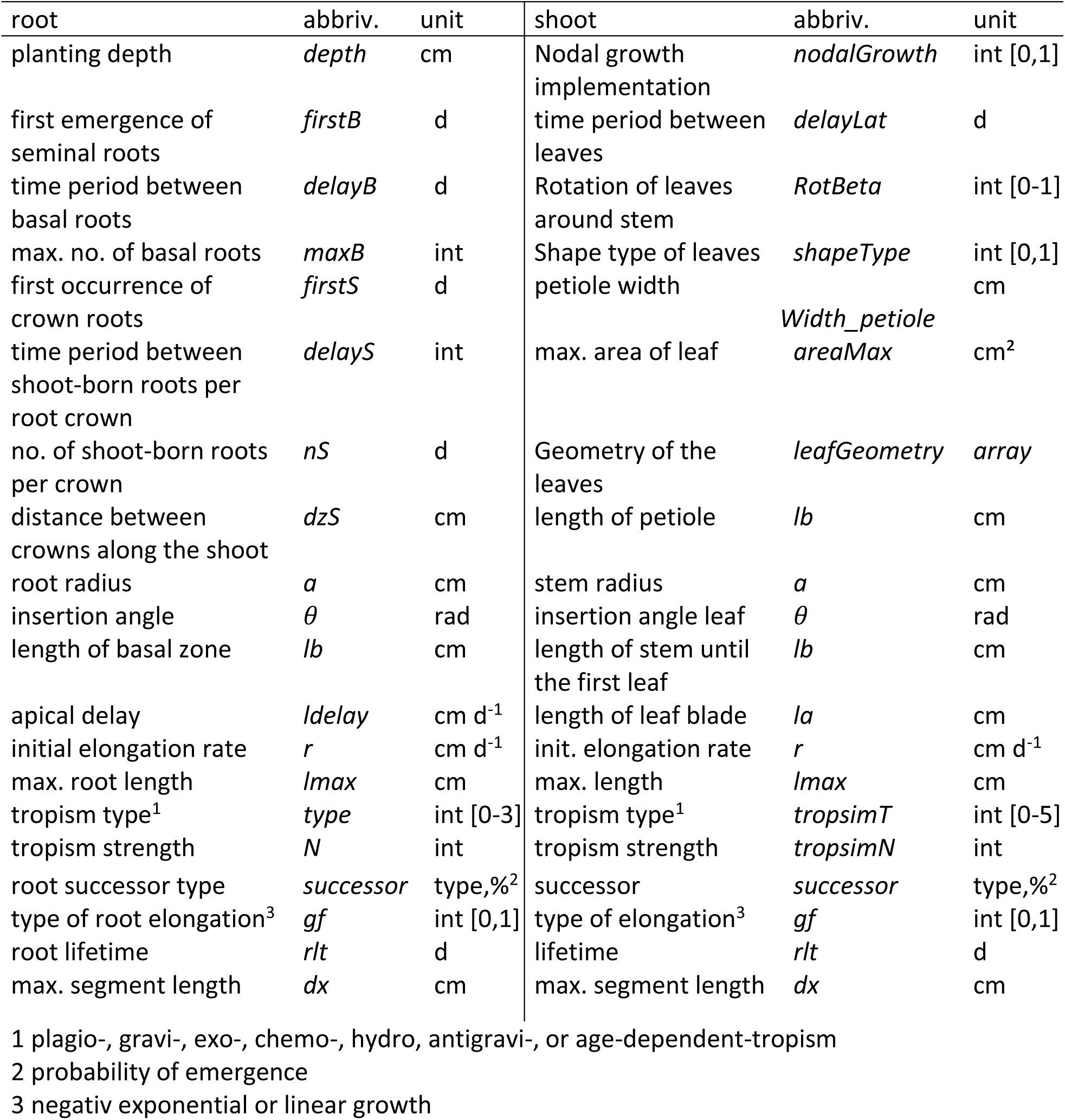
Overview of organ parameters (and their units) that are used to calibrate models with CPlantBox, as used for this study (day: d; integer:int), adapted from Schnepf et al. (2018), Zhou et al. (2020) and Giraud et al. (2023).

**Table 2:**
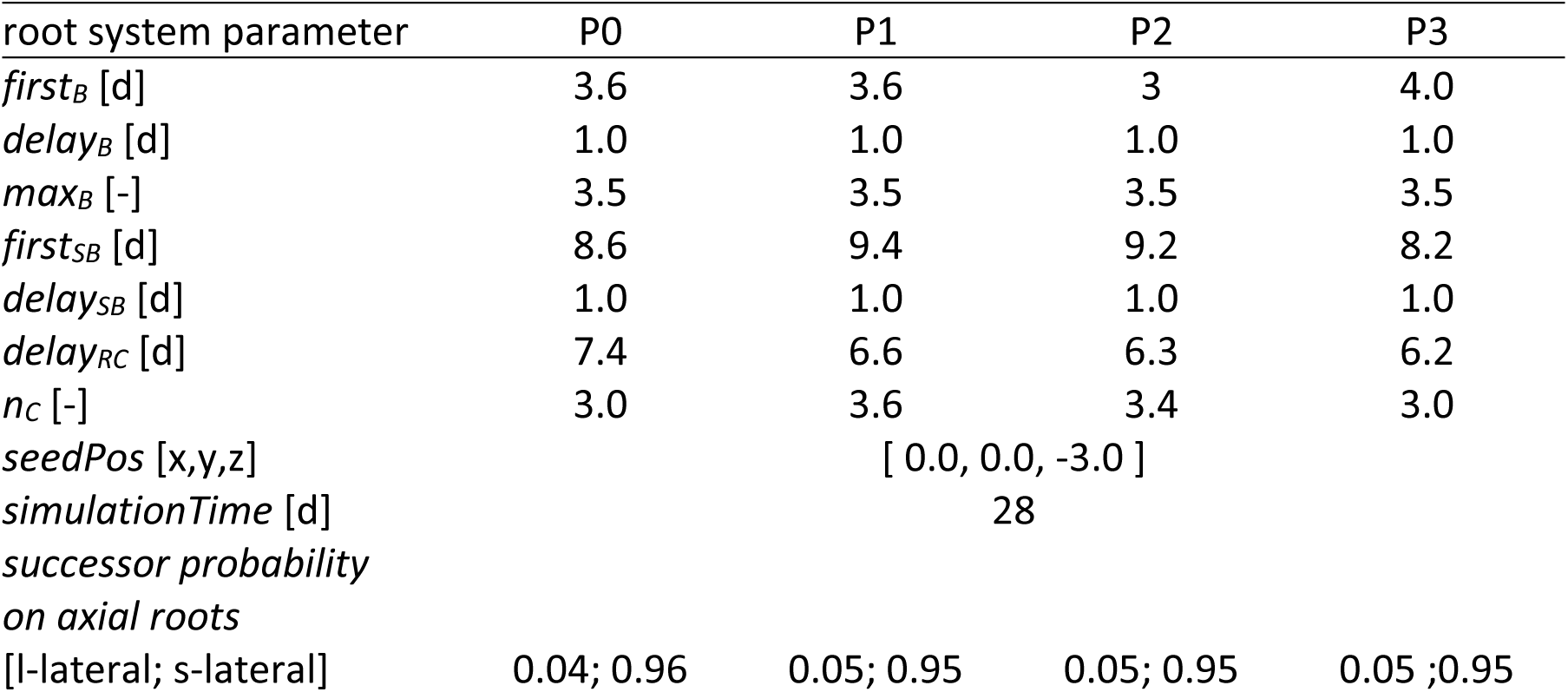
Overview of initial root system architectural parameters for the distinguished P regimes. These parameter describe the initiation time, maximal count and appearance probability of the different lateral root types, the seed position and simulation time.

| root system parameter | P0 | P1 | P2 | P3 |
| --- | --- | --- | --- | --- |
| $first_B$ [d] | 3.6 | 3.6 | 3 | 4.0 |
| $delay_B$ [d] | 1.0 | 1.0 | 1.0 | 1.0 |
| $max_B$ [-] | 3.5 | 3.5 | 3.5 | 3.5 |
| $first_{SB}$ [d] | 8.6 | 9.4 | 9.2 | 8.2 |
| $delay_{SB}$ [d] | 1.0 | 1.0 | 1.0 | 1.0 |
| $delay_{RC}$ [d] | 7.4 | 6.6 | 6.3 | 6.2 |
| $n_C$ [-] | 3.0 | 3.6 | 3.4 | 3.0 |
| $seedPos$ [x,y,z] | [ 0.0, 0.0, -3.0 ] | | | |
| $simulationTime$ [d] | 28 | | | |
| $successor\ probability$<br>*on axial roots* | | | | |
| [l-lateral; s-lateral] | 0.04; 0.96 | 0.05; 0.95 | 0.05; 0.95 | 0.05 ;0.95 |

**Table 3:** Overview of root organ specific architectural CPlantBox parameters for the distinguished P regimes and the parameterset (general) for simulation to evaluate the root system response parameter.

| para- | P level | P0 |  | P1 |  | P2 |  | P3 |  | general |  |
| --- | --- | --- | --- | --- | --- | --- | --- | --- | --- | --- | --- |
| meter | type | mean | sd | mean | sd | mean | sd | mean | sd | mean | sd |
| $a$ | primary | 0.054 | 0.012 | 0.064 | 0.017 | 0.067 | 0.01 | 0.091 | 0.01 | 0.069 | 0.003 |
|  | seminal | 0.052 | 0.011 | 0.062 | 0.016 | 0.066 | 0.01 | 0.081 | 0.011 | 0.065 | 0.003 |
|  | crown | 0.061 | 0.016 | 0.059 | 0.02 | 0.066 | 0.014 | 0.066 | 0.003 | 0.063 | 0.007 |
|  | l-lateral | 0.025 | 0.011 | 0.024 | 0.009 | 0.025 | 0.008 | 0.03 | 0.007 | 0.026 | 0.002 |
|  | s-lateral | 0.025 | 0.007 | 0.028 | 0.01 | 0.025 | 0.008 | 0.04 | 0.012 | 0.030 | 0.002 |
| $l_b$ | primary | 0.8 | 0.899 | 1.879 | 2.385 | 3.183 | 2.236 | 3.777 | 5.681 | 2.410 | 2.033 |
|  | seminal | 2.55 | 2.333 | 3.883 | 2.432 | 3.969 | 4.574 | 1.642 | 0.814 | 3.011 | 1.546 |
|  | crown | 3.161 | 2.454 | 7.216 | 7.564 | 3.473 | 2.487 | 3.924 | 3.025 | 4.444 | 2.468 |
|  | l-lateral | 2.27 | 2.433 | 1.732 | 1.441 | 2.854 | 2.349 | 1.779 | 1.392 | 2.159 | 0.564 |
| $l_{delay}$ | primary | 0.212 | 0.153 | 0.481 | 0.597 | 2.63 | 0.284 | 1.743 | 1.331 | 1.267 | 0.527 |
|  | seminal | 0.499 | 0.364 | 0.941 | 0.784 | 1.038 | 0.864 | 0.94 | 0.535 | 0.855 | 0.230 |
|  | crown | 0.194 | 0.124 | 0.666 | 0.625 | 0.547 | 0.476 | 0.666 | 0.428 | 0.518 | 0.210 |
|  | l-lateral | 0.327 | 0.294 | 0.618 | 0.444 | 0.341 | 0.305 | 0.442 | 0.309 | 0.432 | 0.071 |
| $r$ | primary | 3.951 | 0.766 | 3.35 | 1.252 | 4.417 | 0.865 | 4.627 | 0.486 | 4.086 | 0.317 |
|  | seminal | 3.28 | 1.955 | 2.149 | 1.417 | 2.912 | 0.644 | 3.239 | 1.698 | 2.895 | 0.567 |
|  | crown | 2.981 | 2.693 | 2.556 | 2.902 | 2.29 | 2.146 | 4.886 | 2.583 | 3.178 | 0.319 |
|  | l-lateral | 2.951 | 1.492 | 1.763 | 0.693 | 2.15 | 0.56 | 1.742 | 0.582 | 2.152 | 0.444 |
|  | s-lateral | 2.555 | 2.479 | 5.078 | 4.814 | 5.292 | 4.982 | 5.97 | 5.168 | 4.724 | 1.263 |
| $l_{max}$ | l-lateral | 5.549 | 3.821 | 4.756 | 2.513 | 6.736 | 3.546 | 4.794 | 2.392 | 5.459 | 0.721 |
|  | s-lateral | 1.631 | 1.596 | 1.341 | 1.15 | 0.84 | 0.452 | 1.238 | 1.123 | 1.263 | 0.472 |
| $\theta$ | l-lateral | 1.194 | 0.375 | 1.262 | 0.309 | 1.344 | 0.324 | 1.413 | 0.324 | 1.303 | 0.029 |
|  | s-lateral | 1.194 | 0.375 | 1.37 | 0.346 | 1.396 | 0.327 | 1.413 | 0.324 | 1.343 | 0.023 |
| $l_n$ | primary | 0.466 | 0.045 | 0.457 | 0.085 | 0.536 | 0.122 | 0.545 | 0.187 | 0.501 | 0.060 |
|  | seminal | 0.847 | 0.327 | 0.519 | 0.116 | 0.767 | 0.202 | 0.773 | 0.234 | 0.727 | 0.087 |
|  | crown | 0.847 | 0.327 | 0.628 | 0.244 | 0.811 | 0.419 | 0.754 | 0.124 | 0.760 | 0.125 |
|  | l-lateral | 0.833 | 0.946 | 0.459 | 0.284 | 0.53 | 0.417 | 0.48 | 0.319 | 0.576 | 0.308 |

**Table 4:** Overview of shoot organ specific architectural CPlantBox parameters, as described in Table 1, for the distinguished P regimes.

| para-<br>meter | P level<br>type | P0 |  | P1 |  | P2 |  | P3 |  |
| --- | --- | --- | --- | --- | --- | --- | --- | --- | --- |
|  |  | mean | sd | mean | sd | mean | sd | mean | sd |
| $a$ | stem | 0.187 | 0.01 | 0.16 | 0.02 | 0.166 | 0.014 | 0.13 | 0 |
| $l_n$ | stem | 1.487 | 0.313 | 0.153 | 0.175 | 1.676 | 0.214 | 1.636 | 0.126 |
| $r$ | stem | 0.759 | 0.876 | 0.915 | 1.034 | 1 | 0.772 | 1.129 | 0.66 |
|  | leaves | 7.921 | 2.338 | 7.914 | 2.041 | 8.8 | 2.976 | 10.907 | 3.005 |
| $l_b$ | leaves | 0 | 0 | 0 | 0 | 0 | 0 | 0 | 0 |
| $l_{max}$ | leaves | 38.411 | 7.88 | 42.606 | 14.001 | 52.237 | 17.901 | 49.124 | 15.521 |
| $\theta$ | leaves | 0.705 | 0.306 | 0.773 | 0.055 | 0.794 | 0.403 | 0.739 | 0.262 |
| $delay_{lat}$ | leaves | 3 | 3 | 3 | 3 | | | | |
| $RotBeta$ | leaves | 1 | | 1 | | 1 | | 1 | |
| $Width_{Blade}$ | leaves | 1.638 | | 1.681 | | 1.561 | | 1.563 | |
| $Area_{max}$ | leaves | 54.454 | | 66.695 | | 80.683 | | 71.956 | |

### Root system hydraulics

Based on the created FSPM we calculated *K_rs_*. Our results indicate a close association of P in soil and *K_rs_* (see Figure 2). The *K_rs_* for a root system under high to mild P deficiency was significantly lower than for the root system with a high P supply. After 28 days, the simulated mean *K_rs_* (100 simulations) was between 0.014-0.016 cm^2^d^-1^ for P0, P1 and P2, while P3 reaches a *K_rs_* of 0.021 cm^2^d^-1^ at the same time point. The differentiation in *K_rs_* between the treatments begins between 7 and 10 DAS. Figure 5 shows (A) the temporal evolution of *K_rs_* according to our simulations and (B) in comparison with literature values.

**Figure 5:**
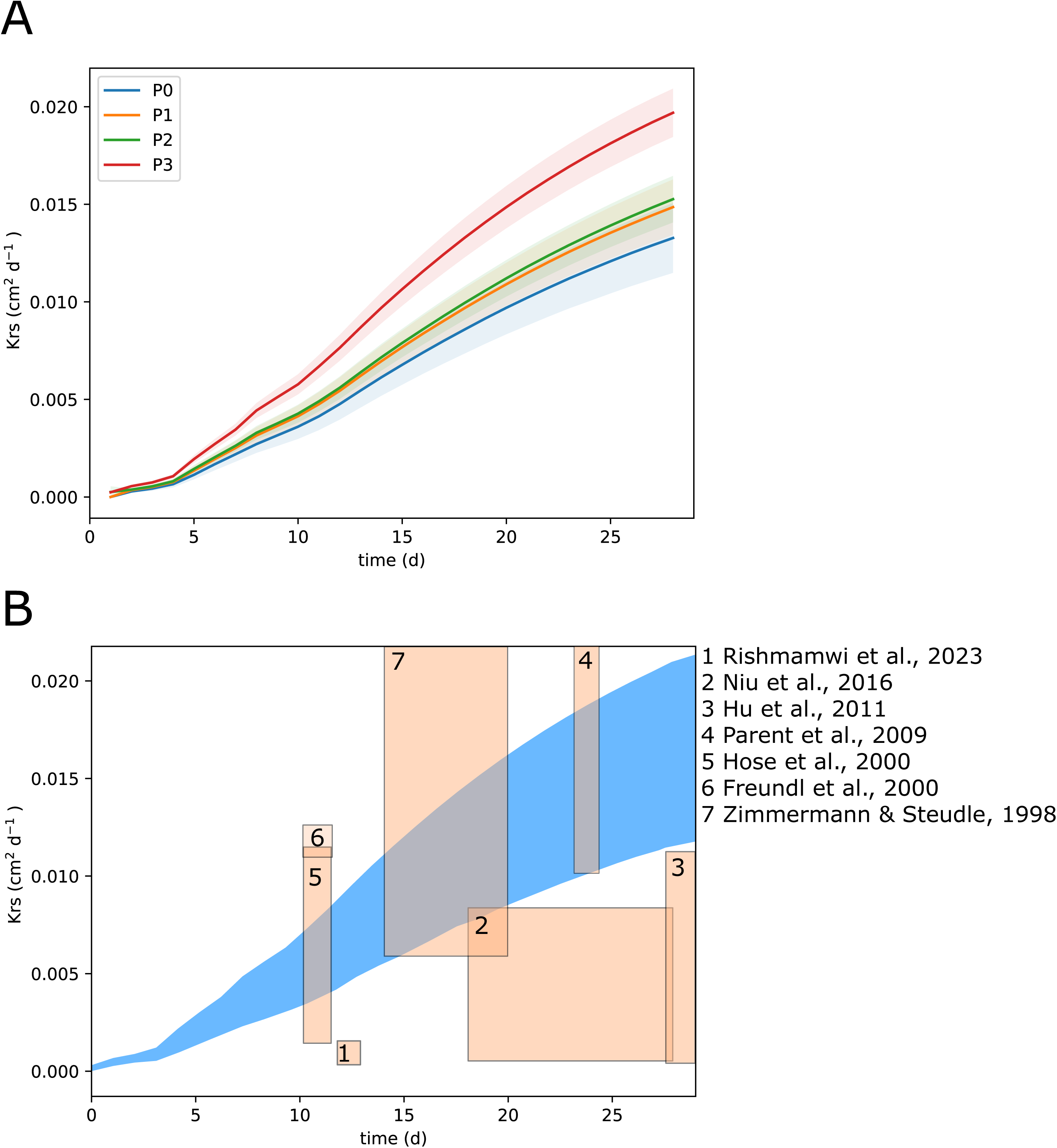
A: *K_rs_* calculated for each P level with 100 simulation runs (shaded areas show the standard deviation to the mean); B:Comparison of different studies investigating *K_rs_* of *Zea mays* with our results (blue).

## Discussion

The here presented study focuses on two main points. First, we conducted a whole plant phenotyping experiment of *Zea mays* cv. B73 under various P availability conditions in rhizotrons to identified which architectural parameters of maize organs are responding most to variations in soil P availability. Second, we parameterized FSPMs based on the previously measured data to understand how root system conductance in maize adapts to the different P availability levels. With additional hydraulic property data (*k_r_*, *K_x_*) from Heymans et al. (2020), which was based on *Zea mays* cv. B73 anatomy, we could calculate the *K_rs_* according to the different structures of the root systems. This allowed us to explore the water uptake capacities of each root system under static soil conditions.

### P levels strongly influence axial root radius and crown root elongation

Having a close look at the architectural parameters of the plant, we could see that initial leaf elongation reacted to P deficiency. We observed clear differences in maximal leaf area depending on the P content in the soil, which originated from significant differences in the initial elongation rate of the leaf between high and low P levels, indicating that the P deficiency was already an important limitation in the initial growing phase of early leaves. Finally, a reduction in the maximal leaf area of the leaves might have disadvantageous effects on water regulation and total photosynthesis. These results might be not surprising, as P deficiency reaction of the plant is mainly linked to a rigid appearance of the shoot (Plénet et al., 2000; Plénet et al., 2000). However, the quantitative empirical values presented here and the derived response functions are valuable additions, as, contrary to most studies, the functions are valid for P levels ranging from strongly limited to sufficient (Lopez et al., 2023).

The influences of P limitation on the root system were more complex to disentangle. Especially since there exist maize genotypes that are considered P efficient and P inefficient. B73 is considered an inefficient line and is thus suitable for investigation on the reaction to P deficiency, since possible reactions might be observed already under mild P stress (Kaeppler et al., 2000). Although it is well known that P deficiency can result in a shallow root system and the promotion of lateral branching (Borch et al., 1999; Zhu and Lynch, 2004; Marschner, 2011), it still remains unclear which adaptions of root system architecture might occur under realistic field conditions linked to real agricultural problems, such as maize cropping on P deficient soils. This is mainly because it is complicated to observe root system responses to P deficiency in field soil. Furthermore, several phenes interact, so phenotypic effects are not always clear to observe in a single organ, although they become more clear from a holistic perspective, when all organs are evaluated together (York et al., 2013). We call this effect a plastic reorganization of the root system. The reorganization effects are complex and our understanding of them is limited (Lynch, 2011). However, modeling approaches have already shown that an increasing amount of seminal roots might be beneficial for P uptake (Perkins and Lynch, 2021), although studies focusing on QTL identification of seminal root count and length report the opposite reaction of *Zea mays* cv.

B73 under lab conditions (Zhu et al., 2006). Our findings do not unequivocally support either of the divergent perspectives reported in the literature. However, the reduced radii of axial roots as response to P deficiency is aligning with previous observations. We could show that there is a high linear relationship between plant available soil P and axial root radii (Sheng et al., 2012; Zhang et al., 2012). Regarding crown root development, we know that a higher number of crown roots is beneficial under P deficiency (Sun et al., 2018). However, past research has indicated that minimizing the amount of crown roots can substantially lower the metabolic expenses associated with root construction, allowing more metabolic energy to be allocated towards root extension (Gao and Lynch, 2016). Following rhizoeconomic paradigm this would suggest that an increased number of crown roots might already result in an initially reduced crown root elongation. Under conditions of nitrogen deficiency, it has been already observed that there is a decrease in the number of crown roots, which is accompanied by an increase in their elongation rate (Saengwilai et al., 2014). For plants under P deficiency the response of crown root elongation is less well-defined. However, our results suggest that crown root elongation in young plants is already an important response parameter for *Zea mays* under P deficiency and has a negative linear response to decreasing P availability in soil.

Overall, the observations in this study are not only meant to investigate shoot and root in terms of biological general validity, but also to parameterize CPlantBox to obtain dynamic FSPMs under various P conditions and to obtain new findings from this model approach. To our knowledge this is the first approach of a detailed whole plant 3D FSPM parameterization of *Zea mays*.

### *Krs* varies between fully fertilized and deficient plants, but not among those with severe to mild P deficiency

*K_rs_* varies due to environmental conditions (Freundl et al., 2000; Hose et al., 2000; Baca Cabrera et al., 2024). It is known that *K_rs_* is influenced by drought (Parent et al., 2009; Hu et al., 2011), osmotic stress (Niu et al., 2016), but also due to genotypic differences (Rishmawi et al., 2023). The *K_rs_* values simulated with our *Zea mays* FSPM are in the same range as the ones given in these studies (7.00×10^-5^ - 2.37×10^-2^ cm^2^d^-1^, see Figure 5B). We found that, when below a specific threshold, soil P deficiency modulates the root system conductance, which might impact young plant vigour. Indeed, the *Zea mays* plants with the highest P supply had a significantly higher *K_rs_* compared with the *K_rs_* for the three lower P supply levels, indicating that, as soon as the plants suffer from P deficiency, the adjustment of the root architecture reduces their water uptake capability. Interestingly, the degree of severity of the P deficiency has no significant influence on *K_rs_*. Changes in *K_rs_* are not solely a consequence of architectural changes, but rather the result of a combination of altered root architectural traits under phosphorus deficiency and the corresponding adjustments in root functional properties that govern water uptake capacity. Especially soil P related changes in root radii and crown root length, due to faster elongation, (as shown by eq. 1&2) influences the root’s radial conductance (eq. 8), which is significantly contributing to observed changes in *K_rs_*.

For the plant’s water uptake that means that on the one hand plants under P deficiency can mitigate drought with water saving strategy, because they have a higher drought stress resilience in beginning drought conditions, since a lower plant water potential is required to maintain uptake and remaining soil water would not be depleted so quickly. On the other hand, higher *K_rs_* could enhance drought recovery following severe drought conditions and would be generally beneficial under sufficient water conditions, since the general capability of water uptake is higher. These are new insights, since studying *K_rs_* experimentally on this high spatio-temporal scale is challenging, due to the complex architecture of root systems, their dynamic interactions with varying soil environments, and the technical difficulties associated with accurately measuring water flow through roots under different conditions (Heymans et al., 2020). With this approach we also showed that computational modeling is overcoming these challenges and could be a tool for improving our understanding of the dynamic modulation of root water uptake mechanisms under P starvation, since all other investigation methods only provide static information at a certain time point and of specific parts of the root system (Shanguan et al., 2005; Mu et al., 2006; Li et al., 2009; Yu et al., 2024).

## Conclusions and Outlook

Our results showed changes of the root system architecture under soil P limitations. Root volume increases linearly with soil P. We identified decreasing axial root radii and crown root elongation as key parameters for root system’s and leaf elongation as main shoot response to P limitation. We combined these results into a functional-structural model to show that maximal potential water uptake capacity does not differ between plants with high and mild P deficiency plants, but between fully P fertilized and P deficient plants. Both, root system anatomy and architecture are key to understanding root system function. Although root system architectural traits, such as volume, do increase linearly to soil P availability, the root system’s capacity to take up water does not follow the same trend. That underscores that root system organization is critical for its function rather than mere total size. To guarantee a better generalizability, it would be important to validate whether these results are applicable in field conditions and across different maize varieties. Further research is required to investigate the effects on older plants. Furthermore, evaluation of how the local intrinsic root hydraulic properties themselves might change under P deficiency and information on the internal P concentration within different plant organs under various P soil conditions would be a valuable addition to the results presented here. This study does not fully account for the complexity and heterogeneity of all soil conditions and cases of extreme P over- or under-supply in natural settings, which can significantly affect nutrient availability and plant growth. While the focus on P is critical, it is important to consider interactions with other nutrients and how they collectively impact plant growth and development. The impact of varying environmental conditions beyond controlled settings on P stress responses is not fully explored and it would be beneficial, if further future studies include a range of genetic diversity within *Zea mays* to understand how different genotypes respond to P deficiency.

## Materials and Methods

### Experimental set-up

We conducted an experiment to gather the necessary dataset, encompassing both shoot and root information over a set time period. Five *Zea mays cv.* B73 plants per treatment were grown in rhizotrons (60 cm × 30 cm × 2 cm) (Pfeifer et al., 2014) under four levels of P availability, thereafter called P0, P1, P2, and P3. The experiment was conducted in a greenhouse at the *Forschungszentrum Jülich GmbH*, Germany (50°54’36”N, 6°24’49”E) from May to June 2022. As substrate, a P deficient luvisol soil from the “Dikopshof” long time fertilization trial (Wesseling, Germany) was used (Schellberg and Hüging, 1997). The initial plant available P concentration in soil (P extracted according to the calcium-acetate-lactate (CAL method)) was 1.8 mg P per 100 g soil (P0). The soil was fully enriched by all other nutrients and sufficiently supplied with demineralized water, so P was the only limiting factor for plant growth. The substrate was additionally fertilized (45% P_2_O_5_, Triplesuperphosphate). The resulting soil P concentration was respectively 3.3 mg 100 g^-1^ for P1, 4.6 mg 100 g^-1^ for P2 and 7.7 mg 100 g^-1^ for P3. Together with P0, these four different P levels represent the different P content classifications for agricultural soils, low B to D range, as proposed by VDLUFA. For each P treatment, five replicates were created. To obtain a high temporal resolution, imaging was first performed daily and, after 3 weeks, every two days. The measurements were performed until 28 days after sowing (DAS).

To phenotype the roots, a daily image of the root system was performed with a “PhotoBox”, equipped with a high resolution camera (Canon Inc., EOS 70D; 14mm APS-C), where the rhizotron was always located at the exact same position, avoiding distortion and image-shift (Pfeifer et al., 2014). This allowed us to take high resolution images of the whole growing root system. During the experiment the rhizotrons are stored in boxes in 45° inclination, so the root system will grow towards and along the window of the rhizotron. The windows remained covered and heat shielded between the measurements, so that the roots were growing in a dark and heat isolated environment. To obtain information about the shoot architecture of the maize plant, we performed a high resolution 2D-RGB measurement with a fixed position horizontally to the plant. The camera (Fujifilm Holdings K.K, X-S10) was equipped with a fixed focal length lens (Fujifilm Holdings K.K, 35mm APS-C). To ensure a good image-processing a uniform blue background was installed. During the measurement the rhizotron was fixed at an angle of 45°, to provide a vertical positioning of the maize shoot. To ensure detailed and accurate data collection, the shoot imaging is conducted just prior to the imaging of the roots. At the end of the experiment a destructive biomass measurement was performed.

### Image processing

The data obtained from root and shoot are available as 2D RGB-images (shoot: JPG, 2080x2080 px; root: JPG, 2268x4862 px). To facilitate the analysis of the images, a mostly automated image processing pipeline was established, streamlining the CPlantBox model parameterization from the experimental data (Figure 6). The first step of image analysis was the segmentation of the targeted organ. The shoot image analysis pipeline started with the segmentation of the maize crop shoot. This was performed by a color-threshold-filter algorithm written in Python based on the OpenCV wrapper PlantCV and the OpenCV library itself (Gehan et al., 2017). The blue background was removed using a color-based filter and only the predominantly green-to-red colored plants were still present after filtering. Then, a semi-automated detection with “Root System Analyzer” was performed (Leitner et al., 2013). The parameters used for CPlantBox were directly derived from “Root System Analyzer”. We used the procedure already successfully applied in Yu et al. (2024). For the root system part, we adapted the method from Bauer et al. (2022) to segment the roots in the image with the a deep neural network model trained with “RootPainter” (Smith et al., 2022). We trained the neural network to ignore small gaps in the root system. However, since some gaps remained we added a feature to “RootPainter”, allowing to manually correct segmentation errors and trace the root by hand if needed. The segmentation results were complete 2D binary root systems. The next processing step was the automated root tracing. “Root System Analyzer” directly provided the input parameter usable for CPlantBox, by manually choosing axial roots and automatically detecting the laterals. Finally, a RSML-file (Root System Marker Language) for every root system and time step was produced by “Root System Analyzer” (Lobet et al., 2015). The first root was always flagged as primary root. To discriminate between crown roots and all other root types, the crown roots were manually flagged in the RSMLs with “Smart Root” (Lobet et al., 2011).

**Figure 6:**
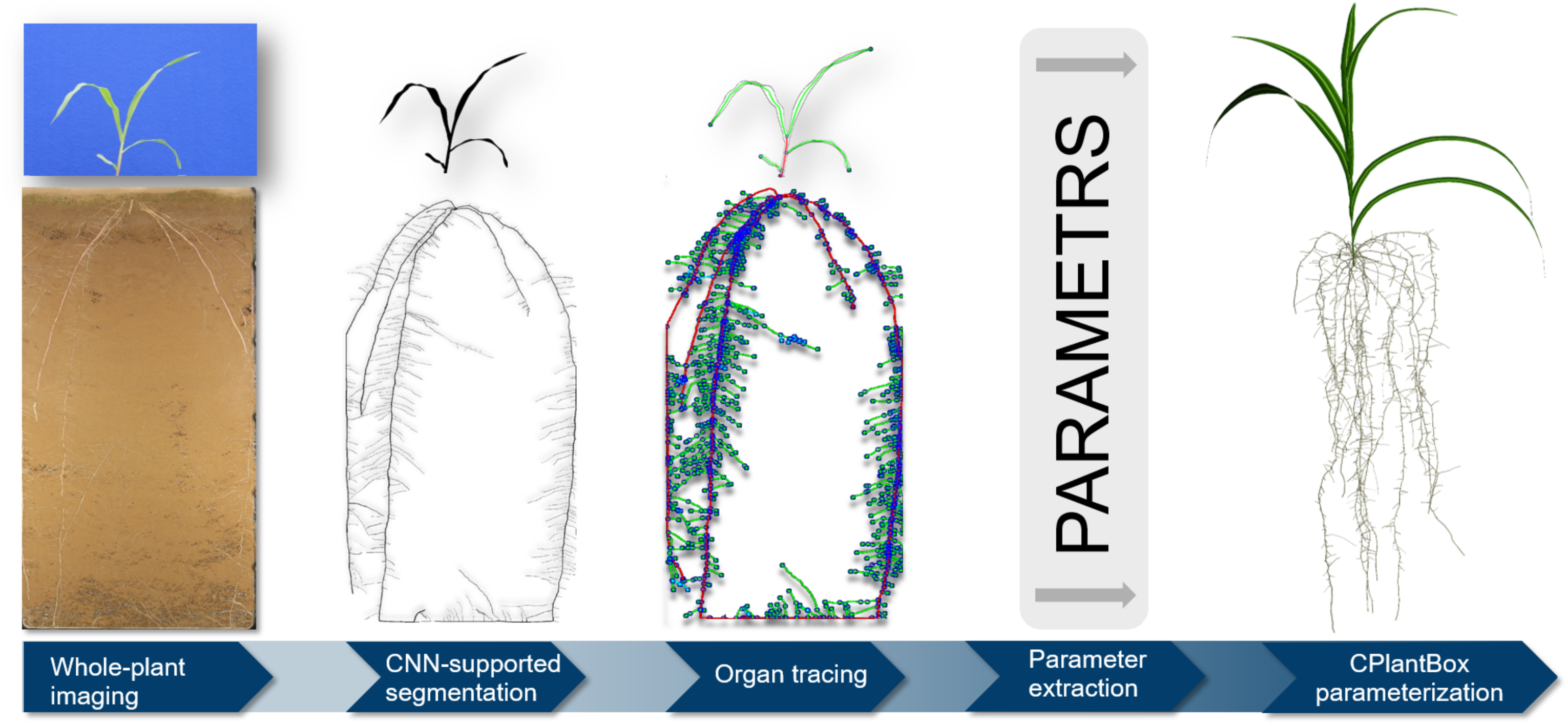
Workflow from experiment to CPlantBox model parameterization.

### CPlantBox parameter extraction

CPlantBox is a modeling platform that can simulate the morphology and 3D topology of the plant, and, among other processes, plant and soil water fluxes (Giraud et al., 2023). To use the CPlantBox modeling framework, plant parameters obtained from real plants are required to create a structure as either a virtual copy of an existing plant or a stochastic variation of a plant, representing the parameterized cultivar, respectively line (Schnepf et al., 2018; Zhou et al., 2020). In terms of plant topology, it is possible to reduce the whole plant architecture to a handful of key-parameters that are the input to calibrate CPlantBox. A precise parameterization of every organ type (e.g, leaf, basal roots) of the shoot and root system is required. This includes plant age at organ emergence, maximal length and initital elongation rate of stem, leaf and every root type. Depending on the organ, initial growth angle, radius, tropism, and branching distance and - pattern have to be defined (Supplementary Figure 2). These parameters were used used as direct model input to simulate the plant structure. We furthermore have parameters that describe general root system traits, such as (first) initiation time, maximal count and appearance probability of different lateral root types and the seed position. We also have organ specific parameters, which had to be measured and calculated for every organ sub-type. Regarding the shoot, this only applied to leaf and stem. For the root system, specific parameter-ensembles were derived for every root-type, respectively primary embryonic root (primary root), seminal roots, crown roots and lateral roots. For maize, there exist also two different types of lateral roots (Heymans et al., 2021). We sub-divided first order laterals into l-(long) laterals, which have branching roots and s-(short) laterals. In a CPlantBox simulation, each parameter is determined using the average value (mean) and variability (standard deviation, sd) from all the data points provided for parameterizing that specific organ. A comprehensive list detailing the parameters, their abbreviations, and the units of measurement can be found in Table 1.

The static root model parameters were directly derived from the RSMLs (Table 1). For the initial elongation rate parameter (*r)* a curve fitting was performed according to eq. 4 (Schnepf et al., 2018). We assumed a maximal root length (*lmax*) of 139 cm from literature and fitted *r* only (Ordóñez et al., 2018). General root system parameters, such as the amount and delay of seminal and crown roots, were evaluated manually from the rhizotrons. For leaves, we also considered negative exponential growth, according to eq. 4. Growth data from leaves that had not yet reached the phase of declining daily elongation rate were not used for the computation of *r*.

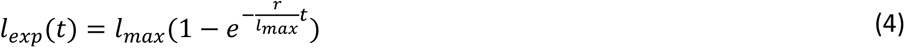

where *t* (*d*) is the time, *lmax* (*cm*) is the maximal length and *r* (*cm d*^−1^) is the initial elongation rate.

### *Krs* calculation

To calculate the root system conductance and assess the water uptake of the plant, information about *k_r_ (d*^−1^) and *K_x_ (cm*^3^ *d*^−1^) are required (Meunier et al., 2018). The root hydraulic properties vary strongly among species, but also among genotypes of the same species (Rishmawi et al., 2023). However, most functional-structural simulations for maize rely on time dynamic hydraulic conductivity profile values from (Doussan et al., 1998), a study conducted 25 years ago that only covers two root types, as highlighted in subsequent studies (Javaux et al., 2008; Postma et al., 2017; Meunier et al., 2020). Beside measuring the radial flow and root anatomy, hydraulic anatomy simulators integrated into new modeling software tools, can assist a more precise estimation these values (Couvreur et al., 2018; Passot et al., 2019; Heymans et al., 2020). This enables new possibilities, such as the hydraulic atlas of *Zea mays* cv. B73 of Heymans et al. (2021). With these parameters, the hydraulic properties of the root system can be defined and the *K_rs_* (*cm*^2^ *d*^−1^) can be calculated according to Couvreur et al. (2012),

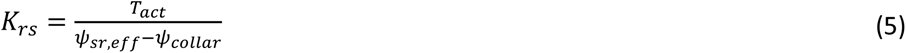

where *ψ*_*sr*,*eff*_ (*cm*) is the *effective* soil-root interface water potential felt by the roots, *ψ*_*collar*_(*cm*) is the plant collar potential and *T*_*act*_ (*cm*^3^ *d*^−1^) is the actual plant transpiration rate and the net sum of the radial water flow rates (*J*_*r*_, *cm*^3^ *d*^−1^) in the roots, since no changes in plant water storage is taken into account. *ψ*_*sr*,*eff*_ is obtained following the method of Couvreur et al. (2012):

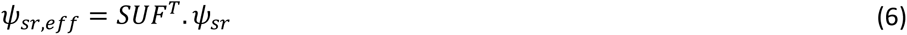

where *SUF* (−) is the vector containing the standard uptake fraction, which is ratio between the water uptake of each root part and the total water uptake of the root system, and *ψ*_*sr*_ the vector of soil water potentials at each root soil interface. *J*_*r*_ is defined as:

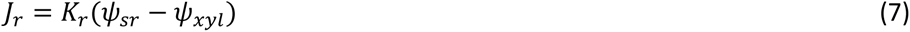

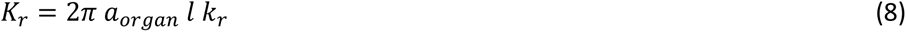

where *K*_*r*_ is the radial conductance (*cm*^2^ *d*^−1^), *a*_*organ*_ is the organ radius, *ψ*_*xyl*_ is the xylem water potential (*cm*), and *l* (*cm*) the length. As we ignore the plant water storage variations, *J*_*r*_ is equal to the changes in axial water flow (*J*_*x*_, *cm*^3^ *d*^−1^) along *l*, so we obtain:

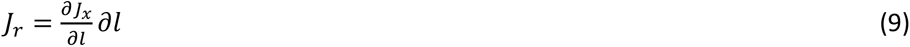

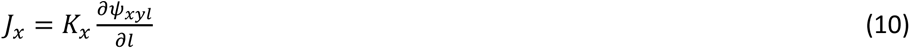

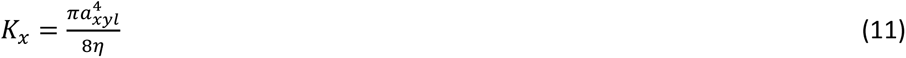

where *a*_*xyl*_ is the xylem radius, *η* (*cm d*^−1^) the dynamic water viscosity, assumed equal to that of pure water at 20°*C*. Eq. 7 - 10 gives us a system of equations that are solved analytically using the method of Meunier et al. (2020), implemented in CPlantBox according to Giraud et al. (2023). The solution yields both *J*_*r*_ and *ψ*_*xyl*_ for a specific set of *k_r_ and K_x_*. Eq. 11 is used for the stem.

We calculated the root hydraulic properties *k_r_* and *K_x_* from the values published in Heymans et al. (2021) for *Zea mays* cv. B73. We assumed that the *k_r_* and *K_x_* did not change between the different P treatments. The changes in radii between P treatments were considered when calculating radial conductance (eq. 8). Although it was shown that the aerenchyma structure can change under P deficiency, the inter-line specific differences in *k_r_* and *K_x_* are much higher in *Zea mays* than the reformation under P deficiency. Furthermore, the aerenchyma reformation of *Zea mays* cv. B73 with a no-P treatment under lab conditions, is reported to be still very moderate (Fan et al., 2007). Finally, the few root hydraulic property data available for maize under P deficiency are hard to use for our model, since they only take a single root type (primary root) into account and are measured for very young plants grown in nutrient solution. The data from Heymans et al. (2021) are given as distance-depending from the root tip distance and for every root type. The conversion from distance-dependent to age-dependent conductivity was done using eq. 12. For a specific distance from the root base *l* (*cm*) the corresponding root segment age (*age*(*l*_exp_), *d*) was calculated with

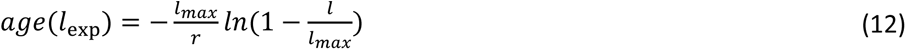

where *r* is the initial elongation rate, obtained from the experiments and eq. 4, *lmax* the maximal root length and *l* is the current measured root length (from experimental data). In contrast to Doussan et al. (1998), we distinguished between primary root, seminal roots, crown roots, l-lateral roots and s-lateral roots. For the parameterization of the shoot organs we followed the simpler approach of (Lobet et al., 2014), where it was assumed that the radial stem conductivity was 0 and the axial stem conductance (*cm*^3^ *d*^−1^) is also calculated according to the Hagen-Poiseuille law (eq.11). We finally calculated the *K_rs_* (eq.5) using the simulated plant architecture and the root hydraulic anatomy based on Heymans et al. (2021).

### Statistical analysis

All statistical analysis, beside the PCA, were performed with Python 3.9.13. For significance testing of the experimentally measured parameter, we applied an ANOVA and Tukey post-hoc test with the “scikit” package (scikit-learn 1.4.2) . All parameters with significant differences (*P*_*value*_ < 0.05) were included in the PCA, namely axial root radii, leaf elongation and crown root elongation and we further added *K_rs_*, root volume, dry matter, P to dry matter ratio and P measured in soil. We clustered for the different P treatments and included the 5 repetitions per treatment. For curve fitting the “scipy” package was used. The PCA was performed with R 4.3.1 (R Core Team 2023) and the “FactoMineR” package. For linear regression models of the identified response parameter, the “sklearn” package was used. Plots were created with the “matplotlib” package.

## Data and code availability

All analysed data, code and model input files used for simulations and to plot the figure are publicly available and released in a GitHub repository https://github.com/Plant-Root-Soil-Interactions-Modelling/CPlantBox/releases/tag/Publication2024 in the folder /experimental/pdef. The image data are available here: doi.org/10.5281/zenodo.11384890. We further transferred the simulation set-up to a docker conatiner for easy access (Supplementary Material 1).

## Acknowledgements

We thank Kerstin Nagel for providing the greenhouse infrastructure and Anna Galinski and Julia Schild for providing assistance with the experiments.

## Author Contributions

FMB conceptualized the study, conducted the experiments, analysed and interpreted the data and drafted the manuscript. DNB implemented the data analysis pipeline, analysed the data and revised the manuscript. MG implemented the model, analysed and interpreted the data and revised the manuscript. JCBC analysed and interpreted the data and revised the manuscript. JV acquired funding, interpreted the data and revised the manuscript. GL and AS acquired funding, supervised the study, interpreted the data and revised the manuscript.

## Supplementary Figures

Supplementary Figure 1: Total volume [*cm*^3^] of simulated root systems with all parameters as measured and with all parameters set as mean and only identified response parameters, elongation of crown roots and axial root radii, set as function of P level in soil.

Supplementary Figure 2: Schematic overview of the different organ parameters required for root and shoot calibration with CPlantBox.

